# Reactivating Memories from a Mathematical Task over a Period of Sleep

**DOI:** 10.1101/2021.10.04.462011

**Authors:** Adrianna M. Bassard, Ken A. Paller

## Abstract

Sleep, especially slow-wave sleep (SWS), has been found to facilitate memory consolidation for many types of learning. Mathematical learning, however, has seldom been examined in this context. Solving multiplication problems involves multiple steps before problems can be mastered or answers memorized, and thus it can depend on both skill learning and fact learning. Here we aimed to test the hypothesis that memory reactivation during sleep contributes to multiplication learning. To do so, we used a technique known as targeted memory reactivation (TMR), or the pairing of newly learned information with specific stimuli that are later presented during sleep. With TMR, specific memories can be reactivated over a period of sleep without disrupting ongoing sleep. We applied TMR during an afternoon nap to reactivate half of the multiplication problems that had previously been practiced. Results showed no effect of TMR on response time or accuracy of multiplication problem solving. Because these results were unexpected, we also used a variation of this paradigm to examine results in subjects who remained awake. Comparisons between the wake and sleep groups showed no difference in response time or accuracy in either the initial test or the final test. Although neither TMR nor sleep differentially influenced multiplication performance, correlational analysis provided some clues about mathematical problem solving and sleep. On the basis of these findings, even though they did not provide convincing support for our hypotheses, we suggest future experiments that could help produce a better understanding of the relevance of sleep and memory reactivation for this type of learning.

## Introduction

Performing well academically can provide many opportunities later in life. Research on the role of sleep in learning, as described in detail below, has produced indications that sleep can be helpful for academic achievement, but many questions regarding the direct impact of sleep on ecologically valid paradigms remain to be addressed. The current experiment focuses on memory reactivation during sleep in the context of mental mathematical performance. Multiplication learning, for example, falls within the general domain of skill learning. When solving multiplication problems, prior knowledge helps you complete them more efficiently and accurately over time. Understanding the effect of sleep on multiplication learning could provide insights into the way we learn that may also extend to other types of skill learning necessary for academic achievement.

A number of studies have explored the role of sleep in academic achievement more broadly. For example, academic performance has been found to be correlated with external factors such as the amount of sleep a student gets as well as the quality of sleep (Taras & Potts-Datema, 2005). In a review of studies exploring the effects of sleep on behavioral measures, Curcio, Ferrara, and De Gennaro (2006) described that a decline in sleep quality and quantity are reflected in increased sleepiness during the day as well as negative impacts on mood, behavior and a decreased ability to concentrate. These changes can negatively affect functions that are necessary for learning and academic success including memory, attention, and problem-solving. Curcio et al. (2006) found a positive correlation between sleep loss and poorer declarative and procedural learning. Restricted sleep also resulted in poorer academic achievement. Furthermore, the benefits of sleep in paradigms requiring mental manipulation have been studied experimentally to some extent. In the case of mental rotation, a night of sleep was found to be more effective in improving performance compared to an equivalent period of wake (Debarnot, Piolino, Baron, & Guillot, 2013). Because sleep has been implicated in academic learning as well as memory retention for a mental task, it is possible that sleep plays an important role in other tasks that require similar processes such as multiplication learning.

The scientific study of memory stability and consolidation over time and with sleep was first introduced by Müller and Pilzecker (1900). This phenomenon was later studied in rodents as researchers found that hippocampal place cells that fired together during spatial tasks were more likely to fire again during slow-wave sleep, or SWS, compared to other active and nonactive cells (Wilson & McNaughton, 1994). These findings support the hypothesis that reactivation during sleep plays an important role for memory consolidation. In an effort to apply these findings in humans, Peigneux and colleagues (2004) measured blood flow in the hippocampus during a wake spatial task and subsequent sleep. Areas found to be active during the wake task were also active during sleep. Furthermore, the amount of hippocampal activity during SWS alone predicted improvement in navigation at a later test (Peigneux et al., 2004).

Whereas the suggestion that hippocampal activity during sleep produces a beneficial effect on memory is consistent with the correlational evidence from Peigneux and colleagues (2004), Rasch and colleagues (2007) sought to more directly test whether memory reactivation during sleep played a causal role in memory improvement. To do so, specific learning episodes were paired with odors that would be later presented during SWS in an attempt to bias the reactivation of memories. In particular, odor cues during SWS led to hippocampal activation and performance improved for the memories associated with the cued odors (Rasch, Büchel, Gais, & Born, 2007).

Rudoy, Voss, Westerberg, and Paller (2009) went further to test whether reactivation could be provoked by auditory cues as well, and whether individual cues could improve individual spatial memories. Altering memory storage by linking odors or sounds with newly acquired information during a learning phase, followed by unobtrusively presentations of the same stimuli during sleep, is a powerful methodology known as Targeted Memory Reactivation, or TMR. Studies in humans have shown that TMR can improve memory in a variety of domains, including spatial, vocabulary, and procedural (Cellini & Capuozzo, 2018; Hu, Cheng, Chiu, & Paller, 2020; Paller & Oudiette, 2018).

One broad domain that is putatively improved after a period of sleep compared to wake is procedural memory, or acquired skills that are refined over time and performed with little conscious awareness. This type of memory can be related to the way multiplication problems are solved. For example, in order to quickly and accurately execute the desired outcome, both procedural and mathematical skills must be fine-tuned with practice over time. In one experiment testing procedural memory, participants learned a number of finger-tapping sequences that corresponded to different melodies (Antony, Gobel, O’hare, Reber, & Paller, 2012). The sequences were each made up of four notes on a keyboard that created a 12-item sequence. After learning, one of the sequences was chosen at random to be presented during SWS. Performance for these sequences was found to improve during a period of wake practice and interestingly improved even more so after a period of sleep with a specific benefit to those melodies that were reactivated (Antony et al., 2012). A recent experiment looked at whether TMR could improve performance in a motor task (Cheng, Che, Tomic, Slutzky, & Paller, 2021). Here, participants used feedback to learn 16 specific arm muscle movements, each paired with a sound, in order to control myoelectric activity. It was determined that reactivation of half of the sounds during SWS improved later performance measured by speed and efficiency (Cheng et al., 2021). Together, these findings demonstrate the ability for TMR during sleep to improve memory in skill learning tasks that involve motor movement.

Just as wake practice can improve performance for procedural memory, practice carried out during wake is important in promoting the skills necessary for computational fluency. This follows from the theory that in order to master number combinations, students must progress through counting and reasoning strategies before these types of problems can be done with fluency (Baroody, 2006). Because sleep and memory reactivation were found to improve procedural memory, it is possible that mathematical learning, which parallels procedural skill learning, would also improve after a period of sleep and TMR.

Multiplication learning can also be related to creative problem solving. Whereas multiplication problems can either be solved by completing a series of simpler calculations or by retrieving the correct answer, problem solving can be done analytically or through insight (Kounios & Beeman, 2009). The analytic approach refers to intentional and conscious strategies which mirrors the way multiplication problems can be solved if broken down into steps. Insight, on the other hand, refers to unconscious computation that allows one to arrive at the answer without conscious knowledge of the steps that were taken to get there (Kounios & Beeman, 2009). Although insight differs from semantic knowledge of multiplication facts, both allow one to arrive at the correct answer suddenly without deliberate strategies. Because strategies for these two types of tasks are similar, understanding the role of sleep for problem solving might provide information in the context of multiplication learning.

Recently, TMR was applied in a problem-solving context (Sanders, Osburn, Paller, & Beeman, 2019). In this experiment, participants were given 2 minutes to work on a puzzle while a 15-second sound clip played. This process was repeated until participants had six unsolved problems. Overnight, half of the sound clips associated with the unsolved puzzles were cued during SWS. The next morning, participants were tested on all of the unsolved puzzles. The ability to solve was found to be greater for those puzzles that were cued overnight compared to those that were not cued. It is proposed that reactivation during sleep may have allowed puzzle concepts to undergo a restructuring process that facilitated later problem solving (Sanders et al., 2019). While this study focused on a variety of difficult problems, it is possible that a similar process occurs when completing multiplication problems. For example, if the multiplication problem sets are computed in one way throughout the experiment before sleep, reactivation during sleep might affect this process allowing for more flexible calculations that could result in performance or response time changes.

Sleep is likely to have many potential influences on later behavior. Sufficient sleep seems necessary for having adequate attentional resources for learning (e.g. Curcio et al., 2006; Taras & Potts-Datema, 2005). In addition, sleep after learning can also be important. In particular, memory reactivation during sleep contributes to the consolidation of memories and may also contribute to problem solving and creativity (Paller, Creery, & Schechtman, 2021). Researchers have used TMR to examine contributions from memory reactivation to skill learning and problem solving but have not used this method to investigate math learning. We sought to take initial steps to fill this gap in knowledge about sleep. For this experiment, it is important to consider how strategies and experience may contribute to math learning. Speed and accuracy improvements in children have been linked to a number of possibilities including: adopting new strategies, efficiently using previously employed strategies, and flexibility in choosing strategies (Lemaire & Siegler, 1995). In an effort to adapt basic number combinations to young adults in the present experiment, more difficult math problems were chosen using 2-digit multiplicands with multipliers 5-9. Additionally, to allow for later improvements in accuracy and response time, participants were allowed 15 seconds to answer each question. We hypothesized that different strategies would likely be implemented depending on the multiplicand and that pairing wake practice with a tone to be played during sleep would selectively benefit the class of problems associated with that sound as a whole for both speed and accuracy. The current study thus allowed us to examine whether reactivation during sleep provides a boost to either accuracy or performance speed.

An initial study was conducted to quantify rate of learning and forgetting in the multiplication learning paradigm. These results allowed us to gauge the amount of practice needed for participants to reach an accuracy level that indicates adequate learning with room for improvement. Then, in Experiment 2 we explored whether targeted memory reactivation during sleep improved performance measured by either accuracy or response time. Finally, in Experiment 3 we tested participants in a variant of the design without intervening sleep.

## Experiment 1

### Methods

#### Participants

Six undergraduate students were recruited and received course credit for participation. Participants entered the lab and completed 30 multiplication problems.

#### Procedure

Problems were presented visually on a screen one at a time and participants were asked to mentally calculate the problem and enter the answer using a keyboard. Correct and incorrect feedback was presented visually and participants pressed the spacebar before continuing on to the next problem. Students completed five continuous blocks of math problems with each block consisting of the 30 problems, as described below. Multiplication problems were displayed one at a time on a screen. Each problem consisted of one of six multiplicands (13,14,16,17,18,19) and one of five multipliers (5-9) for a total of 30 problems. After problems were completed once, they were then all completed again, each time in a different pseudo-random order. Participants were given 15 s to type their response using a keypad and feedback (correct/incorrect) was then presented on the screen. Participants pressed the spacebar when they were ready for the next problem. Each of the 6 problem sets was randomly paired with 1 of 6 unrelated, environmental sounds played each time a problem from this set was presented. After 15 problems were completed, participants were tested on the sound-multiplication class pairs by being asked to type in the correct class after hearing the associated sound. Participants were then asked to generate and answer a problem out loud given the problem set to promote a flexible association between the sound and problem type.

## Results and Discussion

The average response time for correctly answered problems and accuracy for the participants in Experiment 1 was calculated for each block of 30 problems (Figure 1). We determined that participants had the highest level of accuracy in the final block but were able to correctly answer 80% of the problems by Block 2. Response time tended to decrease with each block hitting a plateau near Block 4. Based on these data, we decided participants should complete each of the 30 problems twice to target a level of sufficient practice that also left room for improvement on both measures.

**Figure 1.**
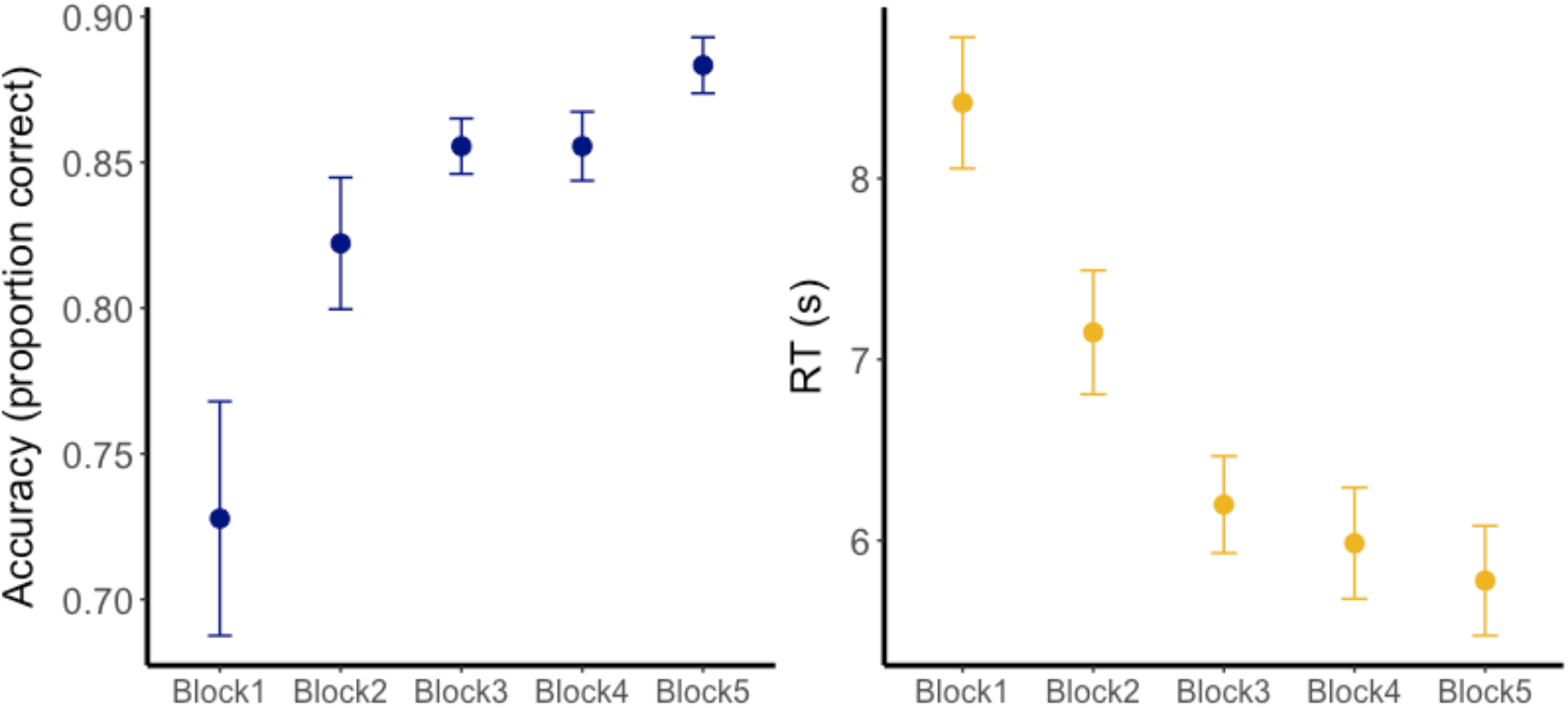
Accuracy (left) and response time (right) learning curves for participants in Experiment 1. Graphs represent the average and standard error of the six participants for each block, which consisted of 30 problems each. Accuracy tended to generally increase with practice while the response time decreased.

## Experiment 2

### Methods

#### Participants

Twenty-one participants (5 males, mean age = 20.62 ± 2.29 (SD)) consented to participate in the experiment and were paid $10 per hour for their time. An additional 6 participants were excluded for failure to receive a minimum of 10 rounds of cues due to arousals from cueing (*n*=5) and for a Pre-test performance below 50% accuracy (*n*=1).

#### Procedure

Participants arrived at the lab between the hours of 11:00 am and 3:00 pm for a 4-hr session. Participants then completed the four stages of the procedure: learning, pre-nap test, sleep, and post-nap test (Figure 1).

##### Learning

Participants completed the same learning procedure described in the pilot experiment with the following exception: the sound test was administered after 10 problems were completed. Three participants began the experiment with three rounds of training and the decision was made to decrease training to two rounds to provide room for improvement for the remaining 18 participants.

##### Pre-nap test

After learning, participants completed each of the 30 problems one time to provide their pre-nap performance. Participants were given 15 s to respond and no feedback was presented on the screen. Again, participants pressed the spacebar between each problem before the next problem was shown.

##### Sleep

EEG recordings were made during sleep using a 21-electrode EEG cap with simultaneous EMG and EOG. EEG, EMG, and EOG data were collected using a 250-Hz sampling rate and filtered with a bandpass from 0.1 to 60.0 Hz. EEG was referenced to the right mastoid. Participants were given 90-120 minutes to nap. Sleep staging was completed offline using criteria of the American Academy of Sleep Medicine with 30-s EEG epochs scored as wake, stage 1, stage 2, stage 3, or REM. SWS is defined as stage 3. Delta power was computed as 0.5-4.0 Hz at electrode Fz.

In order to choose which sounds would be reactivated during the nap, response-time and accuracy for each set was calculated based on the pre-nap test. The problem sets were then separated into two groups, cued and uncued, to minimize the Pre-test response-time and accuracy differences. The auditory cues were presented once the participant reached SWS and stopped if an arousal was detected. If the participant failed to reach SWS within 45 minutes, cues were presented in N2 until an arousal was detected. After the nap, participants were able to remove the cap and gel. This taking an average of 5-10 minutes before returning to the testing room to begin the Post-test.

##### Post-nap test

Each of the original 30 problems, as well as transfer problems were completed. Transfer problems consisted of multiplying the original six multiplicands by the digits 2, 3, 4, and 11 for a total of 24 multiplication problems. Participants were given 15 s to respond and no feedback was presented on the screen. Again, participants pressed the spacebar between each problem before the next problem was shown.

#### Behavioral Measures

A forgetting score (FS) for accuracy was calculated by subtracting the Post-test score from the Pre-test score with a positive FS score indicating more forgotten problems after sleep. For response time, a FS score was also calculated by subtracting performance from the Pre-test from the Post-test with a positive FS score indicating a slower response time after sleep.

The overall cueing benefit for accuracy and response time was calculated by subtracting the FS for cued problems from the FS for uncued problems with a positive score indicating a memory benefit for problems that were cued.

#### EEG and Sleep Analysis

EEG, EMG, and EOG data were collected using a 250-Hz sampling rate and filtered with a bandpass from 0.1 to 60.0 Hz. EEG was referenced to the right mastoid. Sleep staging was completed offline using criteria of the American Academy of Sleep Medicine with 30-s EEG epochs scored as wake, stage 1, stage 2, stage 3, or REM. SWS is defined as stage 3. Delta power was computed as 0.5-4.0 Hz at electrode Fz.

## Results and Discussion

### Behavioral Analysis

Participants showed an average increase in accuracy with practice before sleep. Response time was the slowest during Block 1 of learning and remained consistent between Block 2 and the Pre-test (Figure 3). The learning curves in Figure 2 reflect performance of the 18 participants who completed two rounds of practice before the Pre-test after protocol was changed (described above). In the Pre-test, these subjects had an accuracy of 0.87 ± 0.03 (SE) and response time of 6.41 s ± 0.38 s (SE).

**Figure 2.**
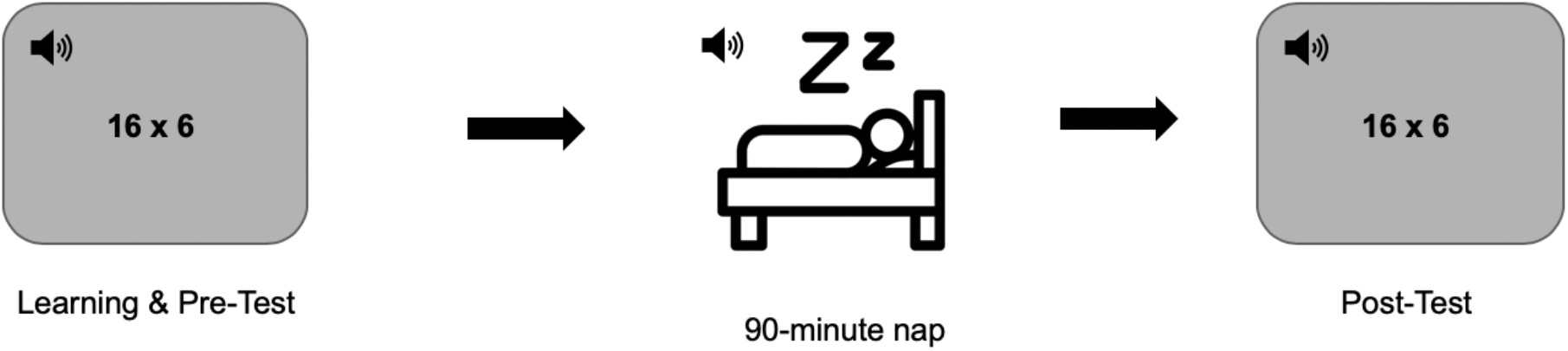
Participants completed learning and the pre-test before taking a 90-minute nap. During stages N2 and N3, three of the six sounds were presented. After waking, participants completed a final Post-test.

**Figure 3.**
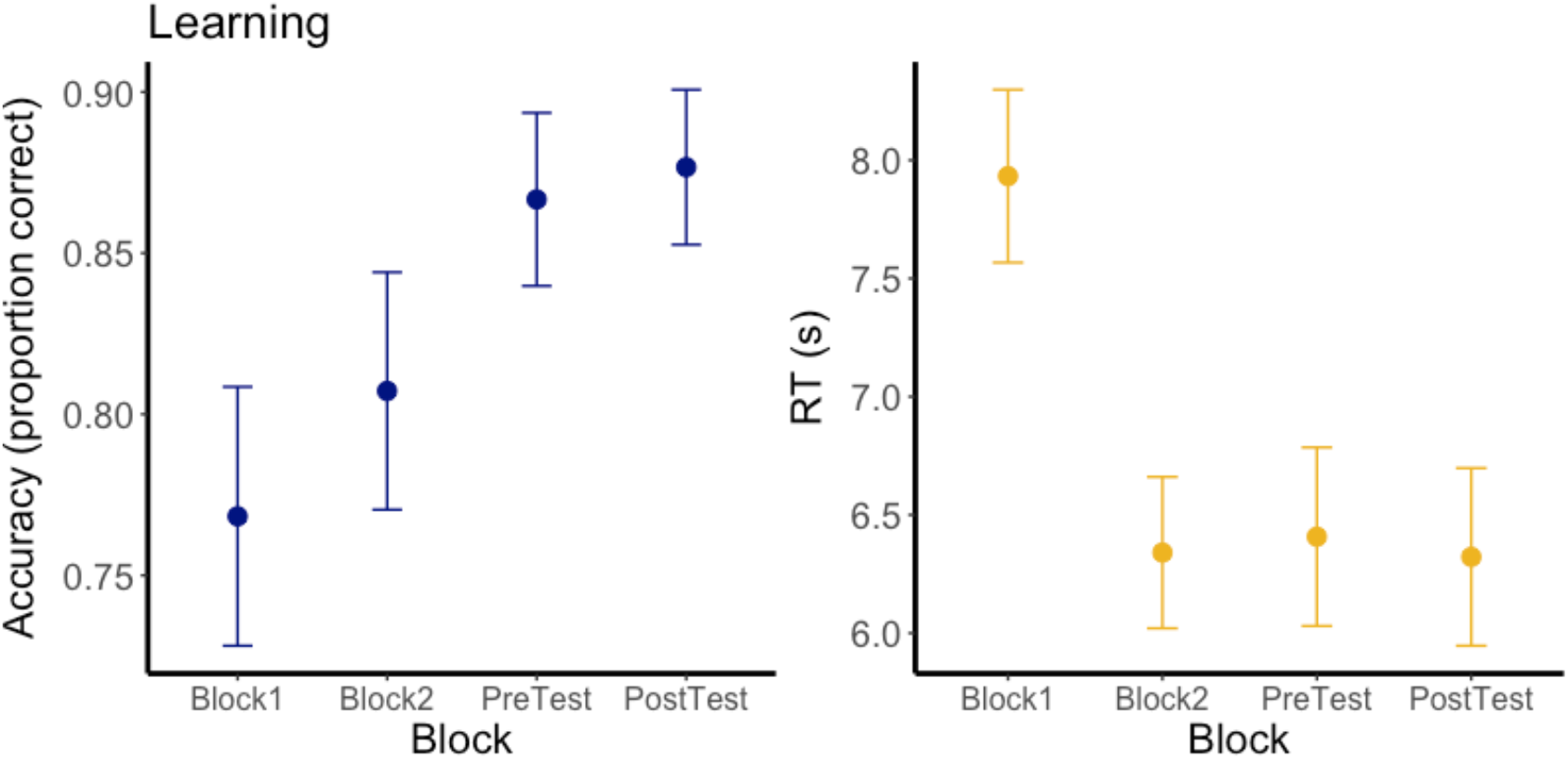
Learning performance for accuracy (left) and response time (right) of 18 participants who completed Block1, Block2, and Pre-test. Each block consisted of 30 problems total.

The final sleep group consisting of all 21 participants had an accuracy of 0.88 ± 0.74 (SE) and 0.88 ± 0.66 (SE) on problems for the Pre-test and Post-test, respectively. The mean reaction time for correctly answered problems was 6.29 s ± 0.33 s (SE) for the Pre-test and 6.32 s ± 0.35s (SE) for the Post-test. A 2 (Test: Pre-test/Post-test) x 2 (TMR: Cued/Uncued) repeated measures ANOVA was conducted to analyze accuracy and to analyze response time. For accuracy there was no main effect of Test (*F*_(1, 20)_ = 0.20, *p* = 0.66), TMR (*F*_(1, 20)_ = 0.24, *p* = 0.63), or interaction of Test and TMR (*F*_(1, 20)_ = 0.68, *p* = 0.42). We also found no main effect of Test (*F*_(1, 20)_ = 0.033, *p* = 0.86), TMR (*F*_(1, 20)_ = 0.42, *p* = 0.52), or interaction (*F*_(1, 20)_ = 0.144, *p* = 0.71) for response time. (Figure 4).

**Figure 4.**
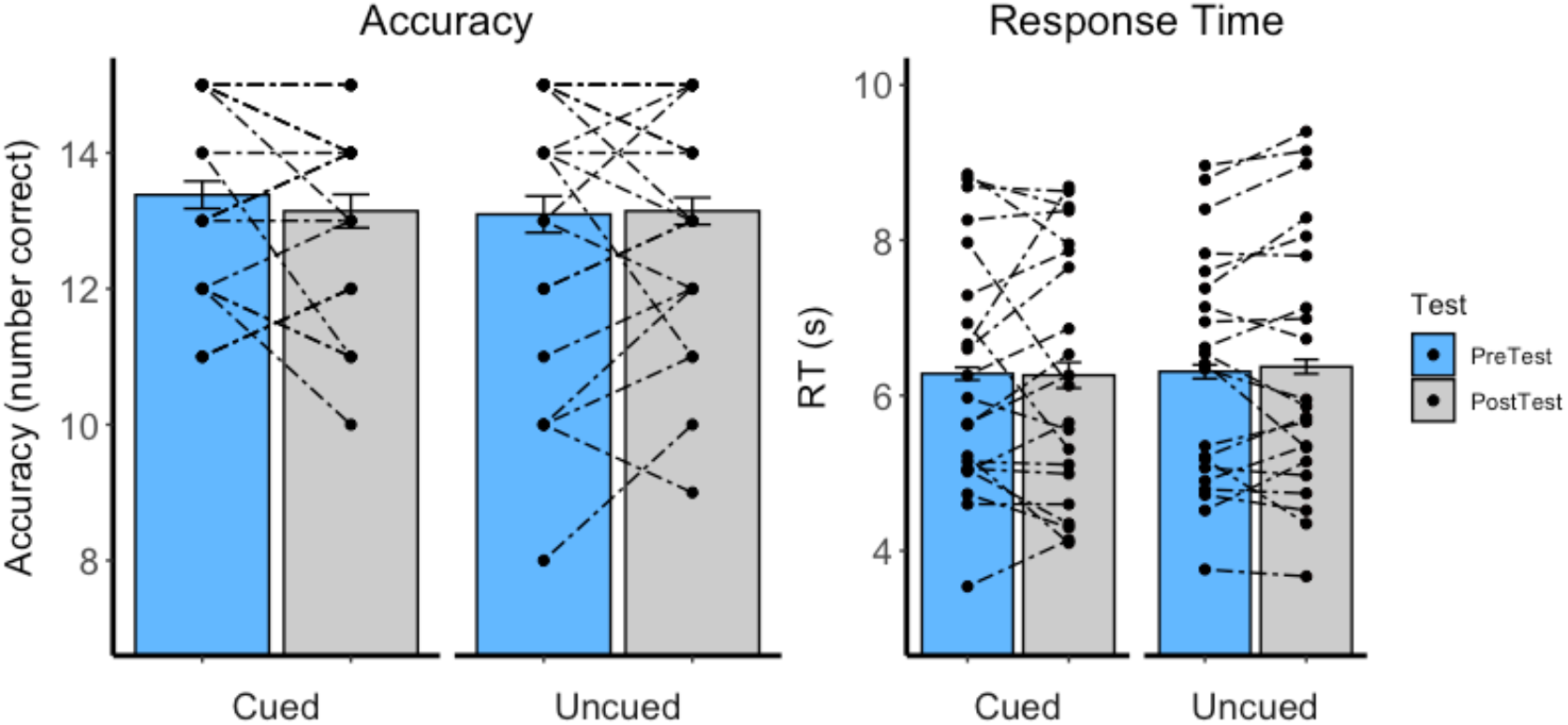
Pre-test and Post-test performance in accuracy (left) and response time (right) for Cued and Uncued sets of multiplication problems.

In order to understand the relationship between Pre-test performance and forgetting, a correlation analysis was performed between these two measures for accuracy and response time. This analysis showed that Pre-test reaction times did not predict FS (*r* (19) = −0.044, *p* = 0.85). Pre-test accuracy, however, was significantly positively correlated with FS (*r* (19) = 0.47, *p* = 0.03) (Figure 5) meaning that a higher accuracy at Pre-test was related to greater forgetting. An explanation for this result might be that participants who perform well on the Pre-test have a greater opportunity to incorrectly answer problems on the Post-test.

**Figure 5.**
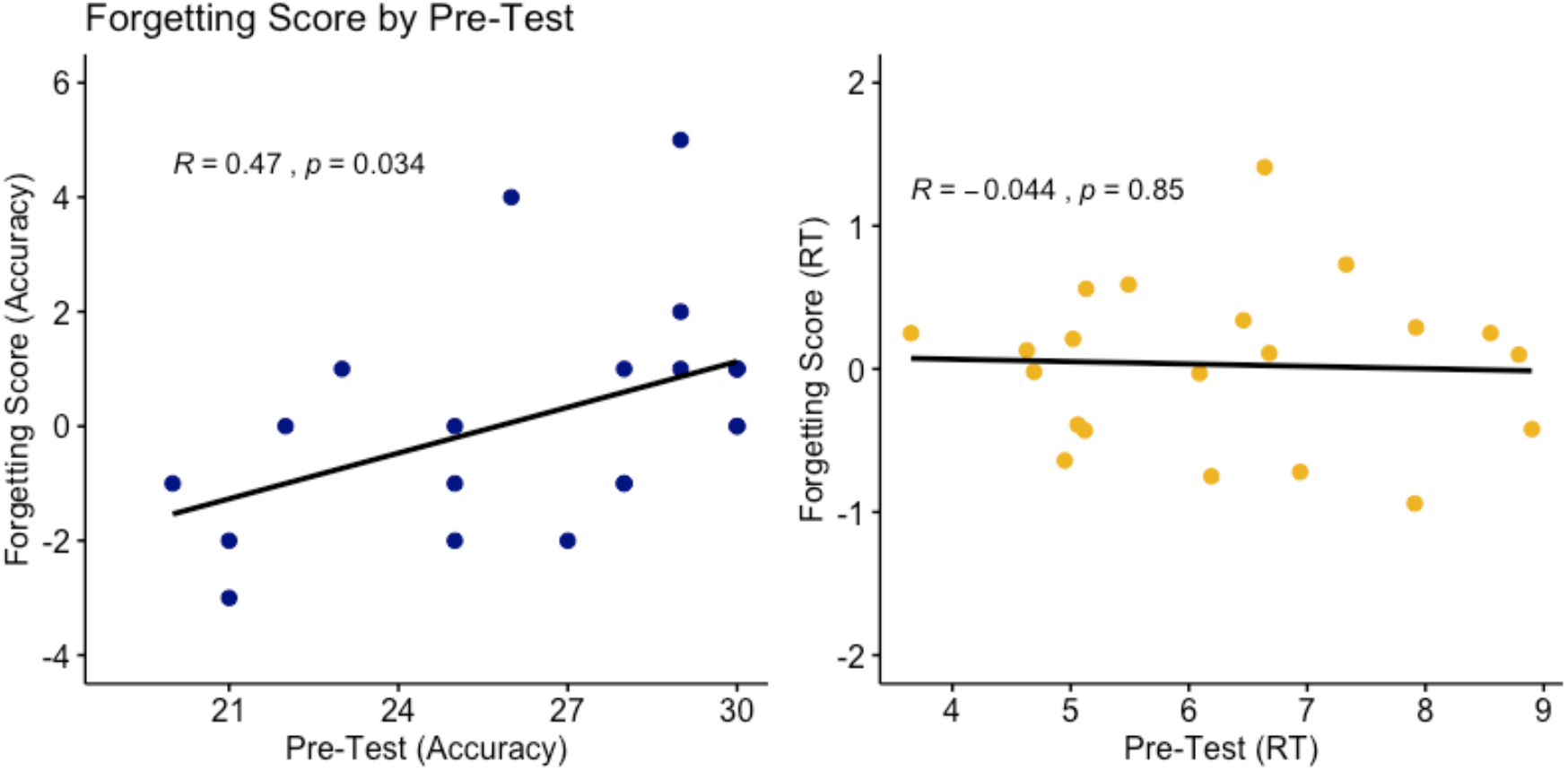
Correlation between Pre-test and Forgetting Score for accuracy (left) and response time (right). Only accuracy showed a significant relationship.

It has been reported that learning that occurs before sleep can also influence the TMR benefit (Creery, Oudiette, Antony, & Paller, 2015). To examine this, we ran a correlation including the pre-sleep performance to predict cueing benefit. We found that the cueing benefit for accuracy was slightly negatively correlated with Pre-test score (Figure 6) while the cueing benefit for response time was positively correlated with Pre-test score (Figure 6) although neither was significant, *r* (19) = −0.13, *p =* 0.58 and *r* (19) = 0.33, *p* = 0.15, respectively.

**Figure 6.**
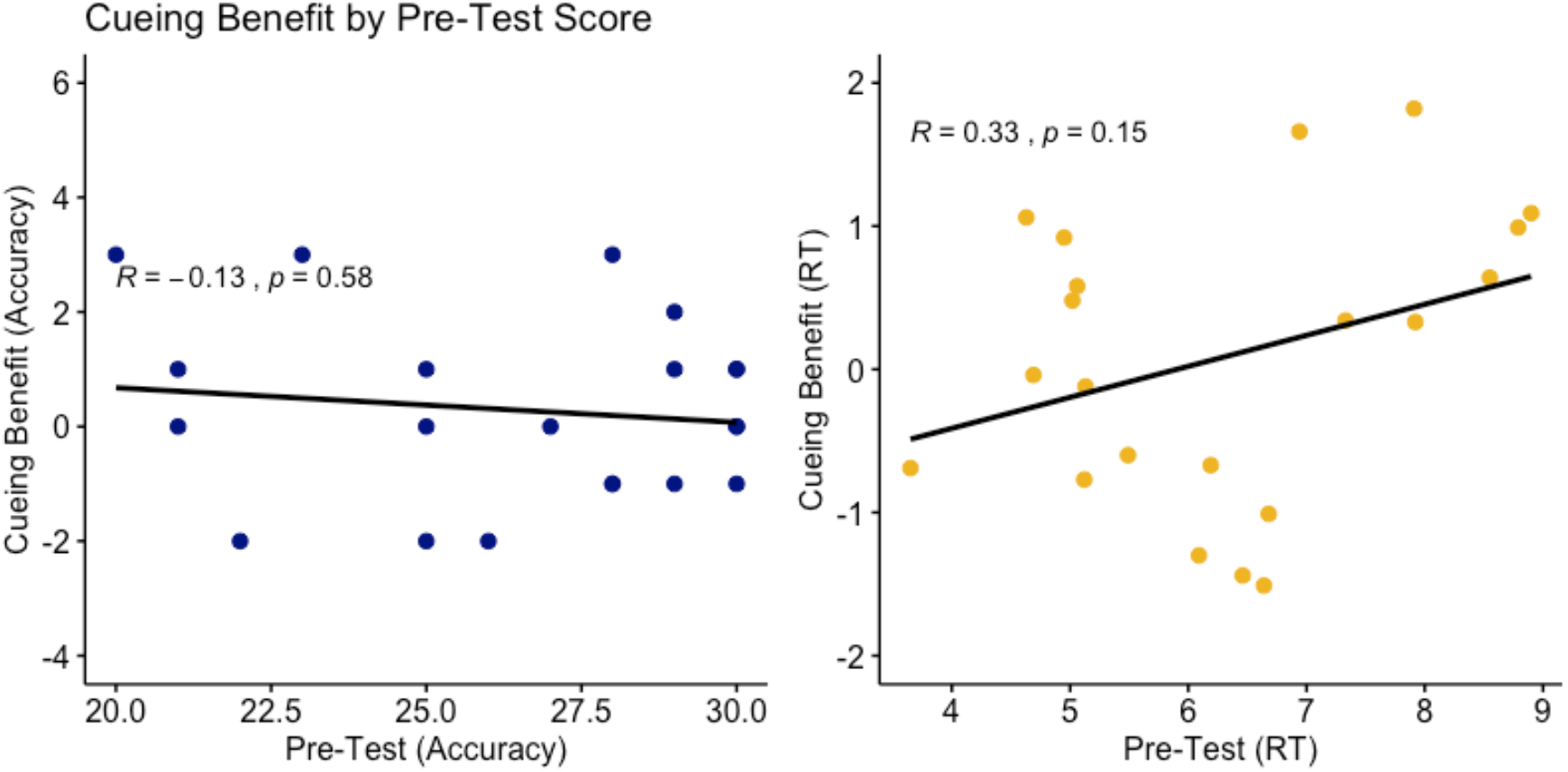
Cueing benefit by forgetting score for accuracy (left) and response time (right). Neither response time or accuracy Pre-test performance is significantly correlated with the cueing benefit.

### Transfer Problems

As described, participants also completed a number of transfer problems in the Post-test. Transfer problems consisted of the original six multiplicands (13,14,16,17,18,19) and four new multipliers (2,3,4,11). Participants had an average accuracy of 0.83 ± 0.03 (SE) overall with an average response time of 6.03s ± 0.28 (SE). For the cued problems, participants had an average accuracy and RT of 0.82 ± 0.03 (SE) and 6.00s ± 0.30s (SE), respectively. For the uncued problems, participants had an average accuracy and RT of 0.83 ± 0.03 (SE) and 6.06s ± 0.32s (SE) (Figure 7). A two-sided *student’s* two sample paired t-test was conducted to determine the effect of TMR:Cued/Uncued on accuracy and response time. It was determined that there was no significant difference on performance for accuracy (*t*(20) =0.44, *p* = 0.67) or response time (*t*(20) = 0.20, *p* = 0.85).

**Figure 7.**
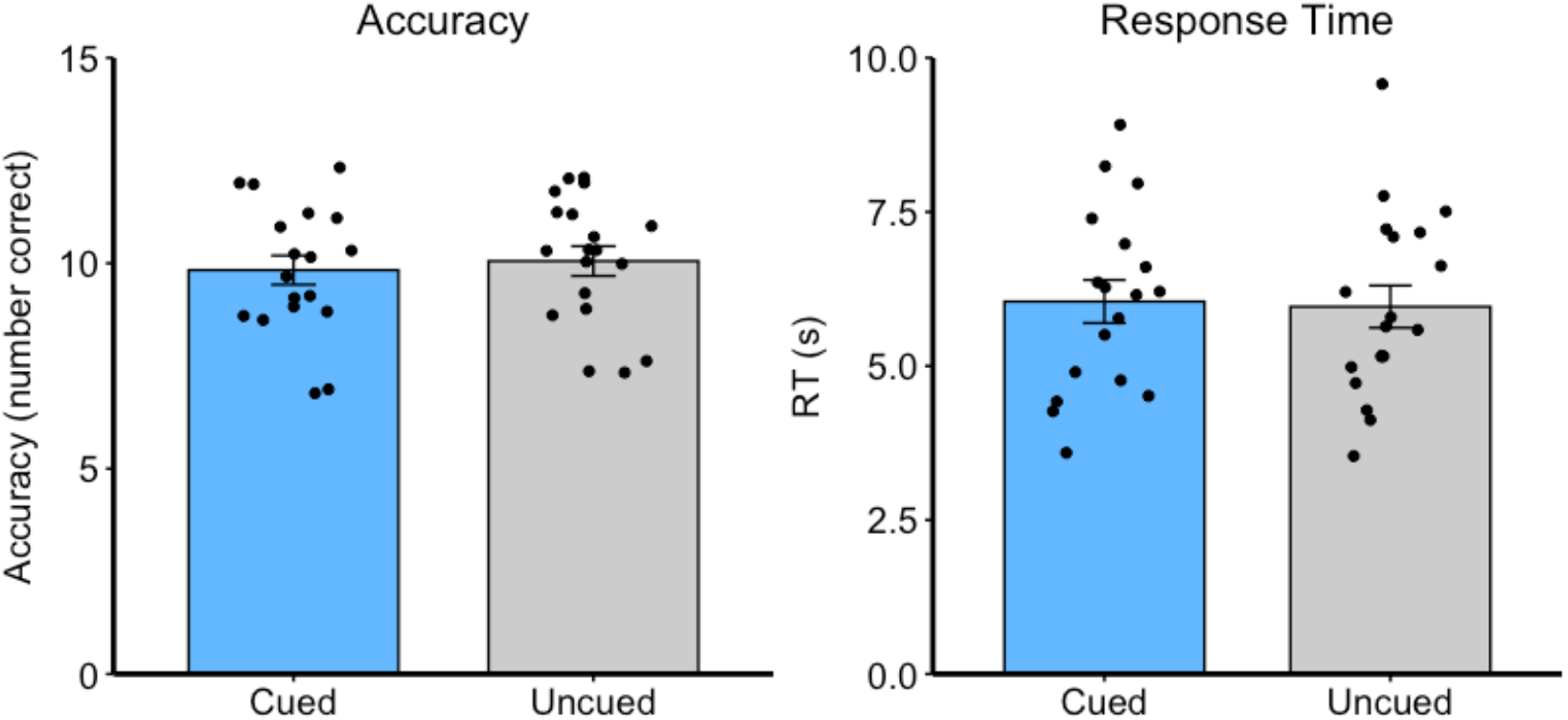
Accuracy (left) and Response Time (right) for transfer problems completed at the Post-test. Performance is separated by cued and uncued problems.

### Item Analysis

Because the multiplication problems may have varied in difficulty based on problem type, we explored average performance for each problem set at Pre-test and Post-test for all subjects including the current and following experiment described in Experiment 2. Looking at this data, performance at Pre-test ranged from 0.82 to 0.95 for accuracy while response time fell roughly between 5-7s. Accuracy had slight changes depending on problem type with Set 13 and 14 having the highest average. Response time, however, revealed larger differences with problem sets 17, 18, and 19 being the slowest on average (Table 2). Based on this data, we ran two 2 (TMR: Cued/Uncued) x 2 (Test: Pre-test/Post-test) repeated measures ANOVA’s for easier problems (Figure 8) and harder problems (Figure 9) for both accuracy and response time. For this analysis, two participants who happened to have cued items fall within either the easy or hard category only were excluded.

**Table 1.** Average time spent in different sleep stages of all 21 participants. Total represents the total time spent in any stage excluding wake. Time cued represents the average time a participant was cued during the nap.

| Sleep Stage | Time (min) ± SD |
| --- | --- |
| N1 | 25.38 ± 19.78 |
| N2 | 32.81 ± 16.36 |
| N3 | 23.48 ± 14.25 |
| REM | 0.1 ± 0.44 |
| Total Sleep Time | 76.48 ± 16.79 |
| Time Cued | 14.45 ± 8.15 |
| Sleep Disruption Index Score | 0.14 ± 0.09 |

**Table 2.** Results by Multiplicand Set (Mean ± SE)

|  | **Pre-test Accuracy** | **Pre-test RT** | **Post-test Accuracy** | **Post-test RT** |
| --- | --- | --- | --- | --- |
| Set 13 | 0.95 ± 0.02 | 5.63 ± 0.27 | 0.94 ± 0.02 | 5.37 ± 0.22 |
| Set 14 | 0.95 ± 0.01 | 5.82 ± 0.29 | 0.91 ± 0.02 | 5.72 ± 0.27 |
| Set 16 | 0.88 ± 0.03 | 5.76 ± 0.27 | 0.87 ± 0.02 | 5.78 ± 0.25 |
| Set 17 | 0.87 ± 0.03 | 6.46 ± 0.33 | 0.90 ± 0.02 | 6.42 ± 0.35 |
| Set 18 | 0.88 ± 0.03 | 6.27 ± 0.30 | 0.86 ± 0.03 | 6.20 ± 0.31 |
| Set 19 | 0.82 ± 0.04 | 6.68 ± 0.31 | 0.84 ± 0.03 | 6.79 ± 0.33 |

**Figure 8.**
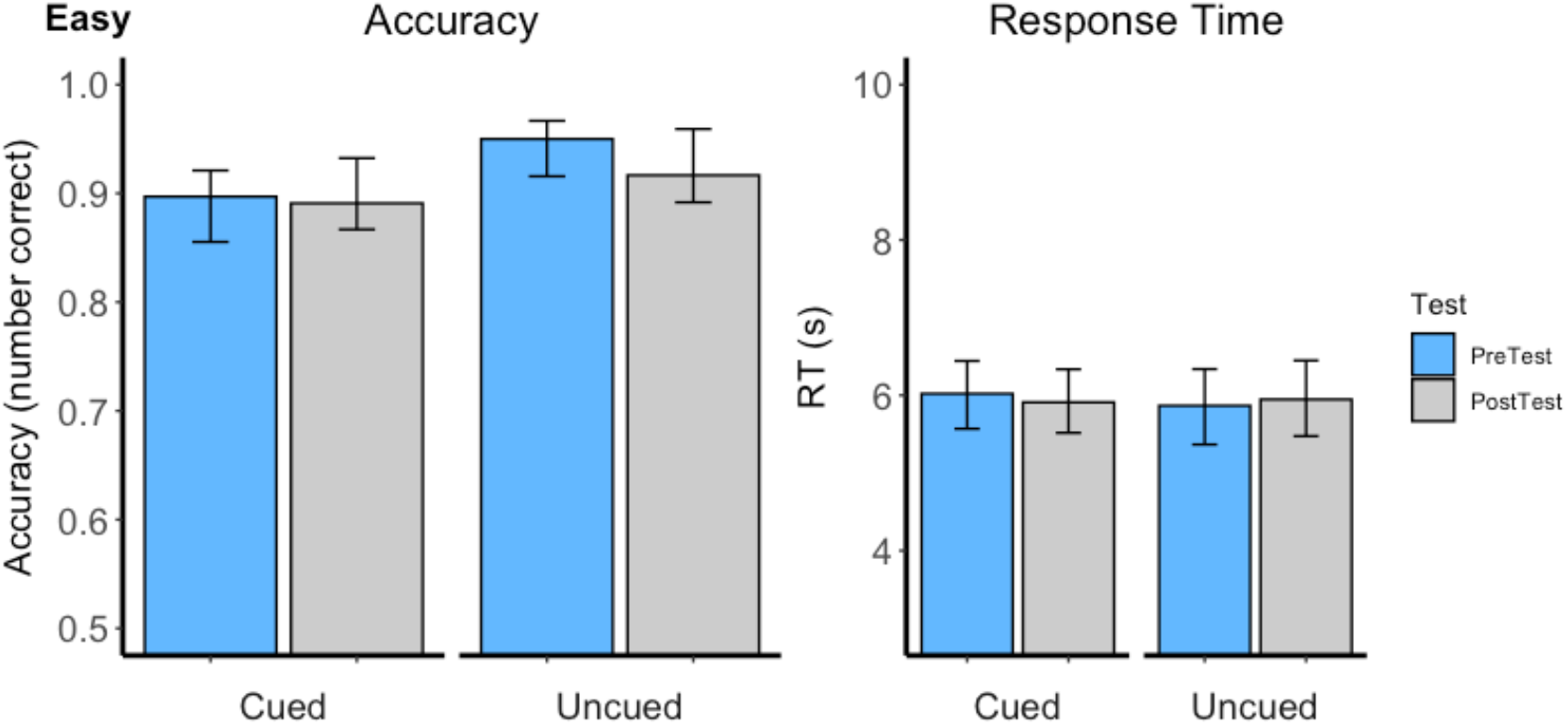
Performance for cued and uncued problem sets 13, 14, and 16.

**Figure 9.**
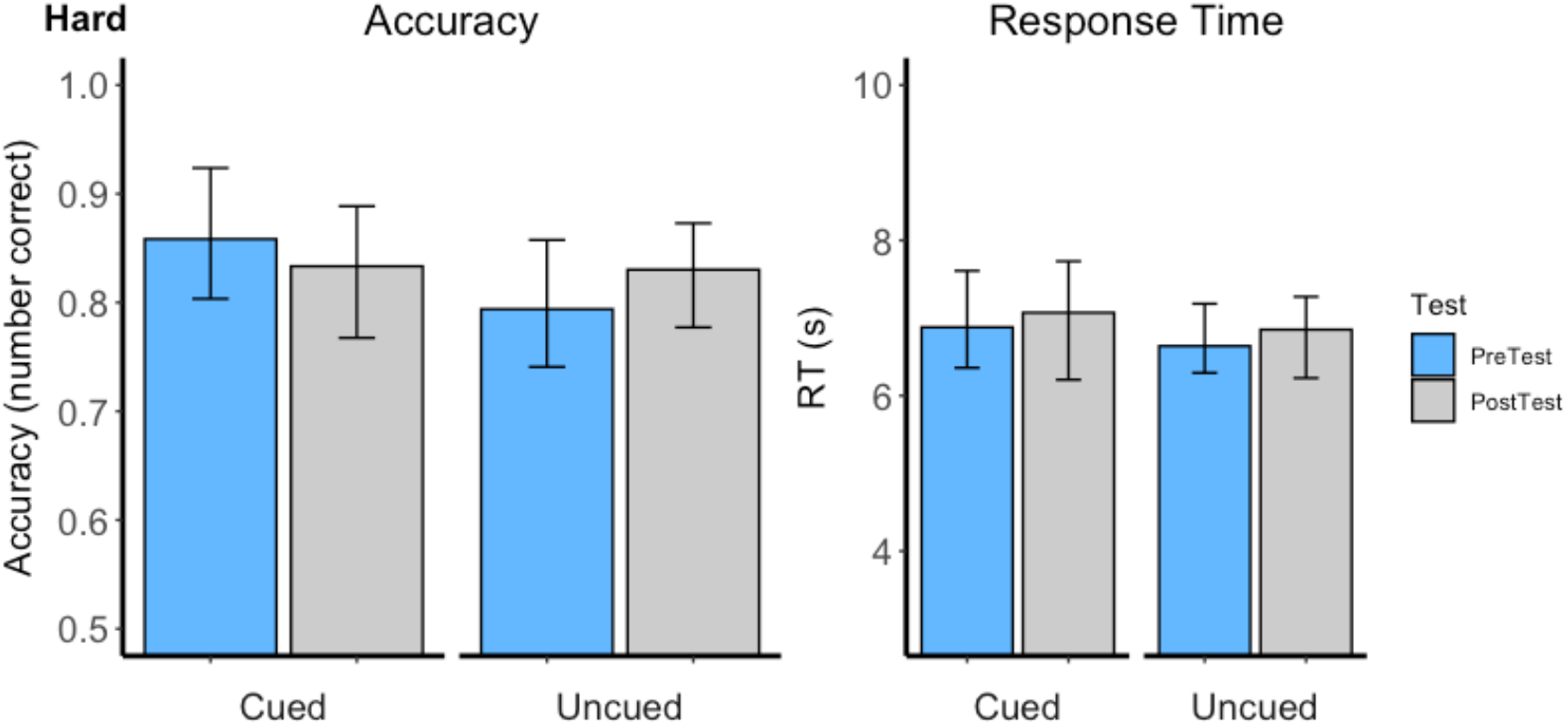
Performance for cued and uncued problem sets 17, 18, and 19.

For easy problems (sets 13, 14, and 16), when looking at accuracy, there was no main effect of Test (F_(1,17)_ = 0.78, *p* = 0.39, TMR (F_(1,17)_ = 0.34, *p* = 0.57) or Test x TMR interaction (F_(1,17)_ = 0.78, *p* = 0.39). Similarly when looking at response time, there was no main effect of Test (F_(1,17)_ = 0.021, *p* = 0.89), TMR (F_1,17)_ = 1.43, *p* = 0.25), or Test x TMR interaction (F_1,17)_ = 0.01 *p* = 0.93). The same analyses were ran looking specifically at the more difficult problems (sets 17, 18, and 19). For these problems, there was no main effect of Test (F_(1,17)_ = 0.10, *p* = 0.76), TMR (F_1,17)_ = 0.32, *p* = 0.58), or Test x TMR interaction (F_(1,17)_ = 0.85, *p* = 0.37) for accuracy. For response time, there was also no main effect of TMR (F_(1,17)_ = 3.07, *p* = 0.08), Test (F_(1,17)_ = 0.77, *p* = 0.39), or Test x TMR interaction (F_(1,17)_ = 0.00, *p* = 0.99).

### Sleep Analysis

Table 1 shows the average time spent in each sleep stage. Participants spent the majority of the nap in N2 with similar amounts in stages N1 and N3. Few participants reached REM in the 90-minute nap.

We first analyzed relationships between performance measures and different measures related to sleep. For behavior, we continued to analyze the cueing benefit and the change in performance measured before and after sleep for both accuracy and response time. Because SWS is typically considered crucial for memory reactivation, we looked specifically at the time spent in Stage 3 and delta power over the nap. When looking at time spent in N3, we found that there was no significant relationship with cueing benefit (Figure 10AB) or amount forgotten (Figure 10CD). We next examined the relationship between the same behavioral variables and delta power. Here we found no significant relationships between delta power and accuracy or response time.

**Figure 10.**
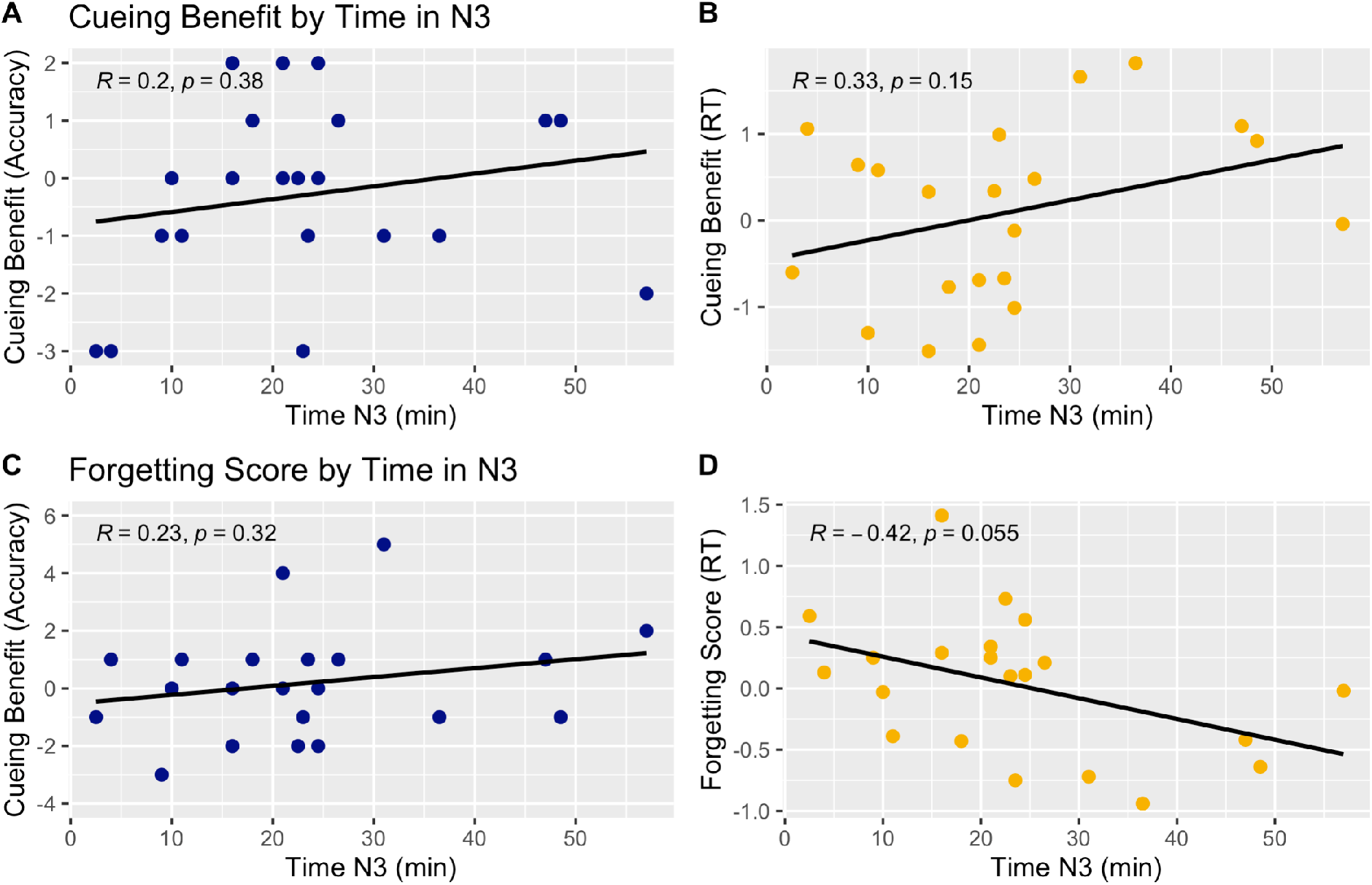
Correlation between the cueing benefit and time spent in N3 over the nap for accuracy (**A**) and response time (**B**). Correlation between forgetting score and time in N3 for accuracy (**C**) and response time (**D**). Time in N3 is not significantly related to cueing benefit or FS.

Although some of the described correlations did not reach significance, there are consistent trends found relating to the behavioral measure of interest. When looking at cueing benefit, we found that time in N3 and delta power both have a positive relationship with response time and accuracy. Conversely, for forgetting scores, time in N3 and delta power are negatively correlated with response time but positively correlated with accuracy.

For the next sleep analysis, we analyzed the EEG response to the cues that were presented during the nap. It was previously reported that participants who reported sleep disruption as a result of TMR cues did not have a memory benefit compared to those who were not disturbed (Göldi & Rasch, 2019). To assess sleep disruption, the absolute change in EEG power at Cz was added across the frequency band from 0.38 to 20.35 during the 5 seconds after cue onset relative to the 5.0 seconds before the cue as described by Whitmore, Bassard, and Paller (2021). We then tested the correlation between this measure of sleep disruption and memory benefit for cued items. Based on this analysis, we concluded that sleep disruption following the cues did not influence the cueing benefit for accuracy (*r* (19) = −0.18, *p* = 0.44) or response time (*r* (19) = 0.10, *p* = 0.66) (Figure 12).

**Figure 11.**
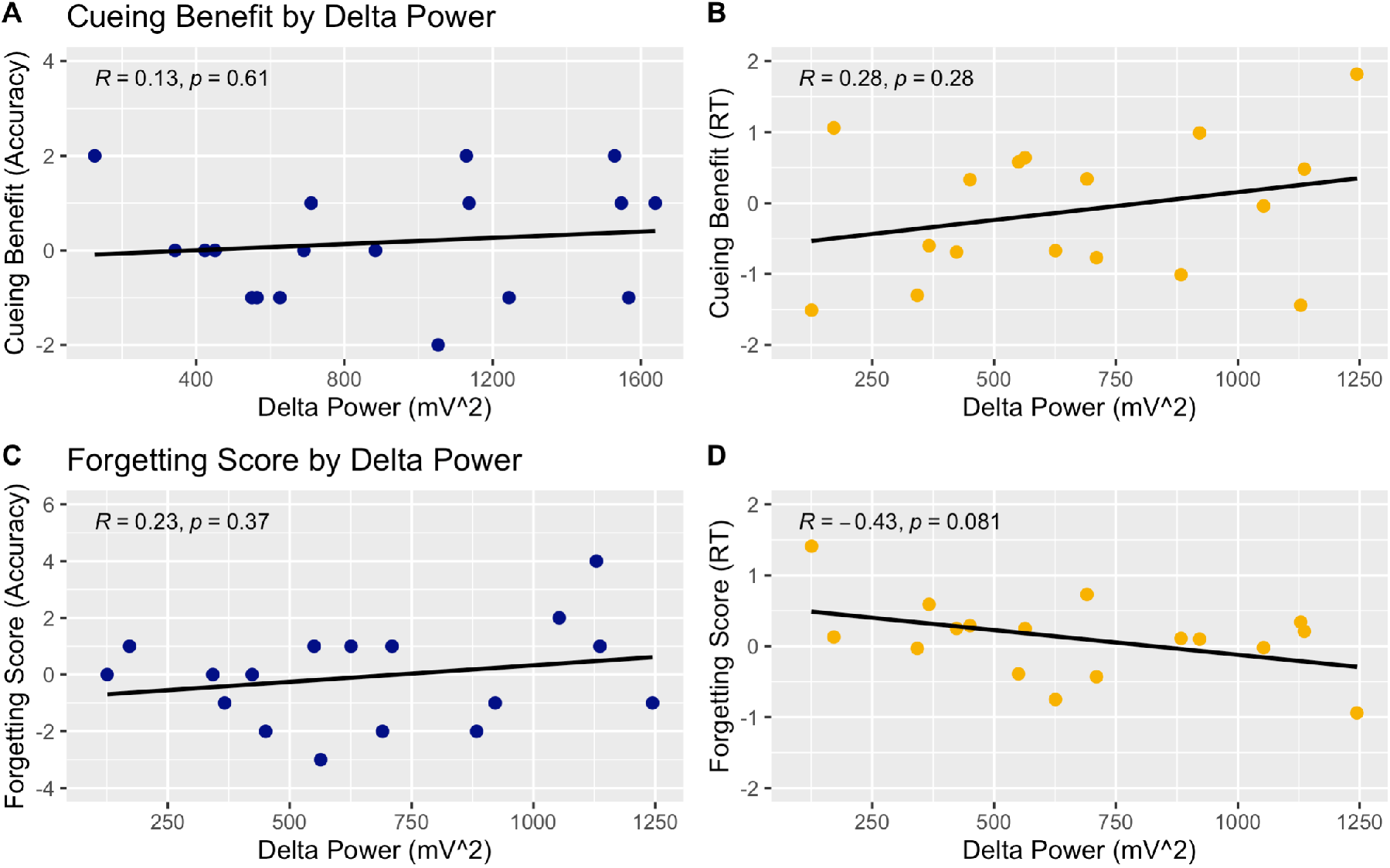
Correlation between the cueing benefit and delta power over the nap for accuracy (**A**) and response time (**B**). Correlation between the forgetting score and delta power for accuracy (**C**) and response time (**D**). Delta power is not significantly correlated with the forgetting score.

**Figure 12.**
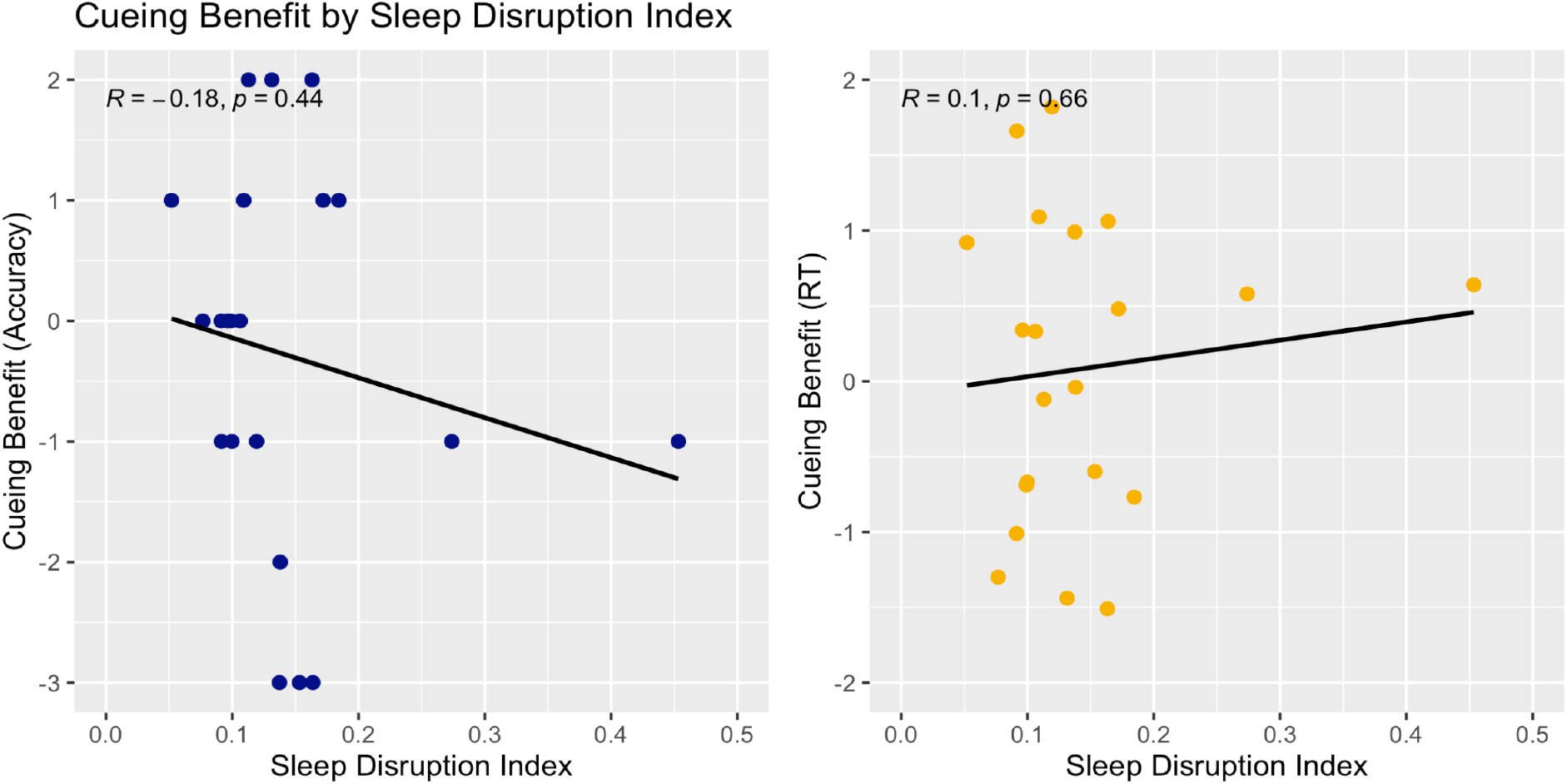
Correlation between the sleep disruption index score with the cueing benefit for accuracy (left) and response time (right). Sleep disruption was not significantly related to the cueing benefit for either measure.

In the the previous exploration of sleep disruption, participants had sleep disruption index scores falling between 0.08 and 0. 28 and were able to be classified as having either low or high sleep disruption (Whitmore et al., 2021). The majority of participants in this study fell within this range aside from one participant who scored above 0.4. These participants, however, did not have scores that could be stratified in this way to explain cueing benefit.

Finally, we ran a multiple regression to predict the cueing benefit for accuracy and response time. Predictors for each included the amount of time spent in N3, delta power, and the calculated sleep disruption index value. We first examined the correlation matrix between the three predictors to assess multicollinearity (Figure 13). Sleep disruption had a correlation value of −0.29 with delta power and −0.40 with minutes in N3. Unsurprisingly, the time spent in N3 was had a positive correlation value of 0.76 with the predictor delta power. The multiple regression analysis revealed that the overall model including minutes in N3, sleep disruption index, and delta power was not significant in predicting the cueing benefit for accuracy (R^2^ADJUSTED (17) = 0.05, *p* = 0.57) or response time (R^2^ADJUSTED (17) = 0.15, *p* = 0.13 with all individual predictors being nonsignificant as well (all *p*’s > 0.10).

**Figure 13.**
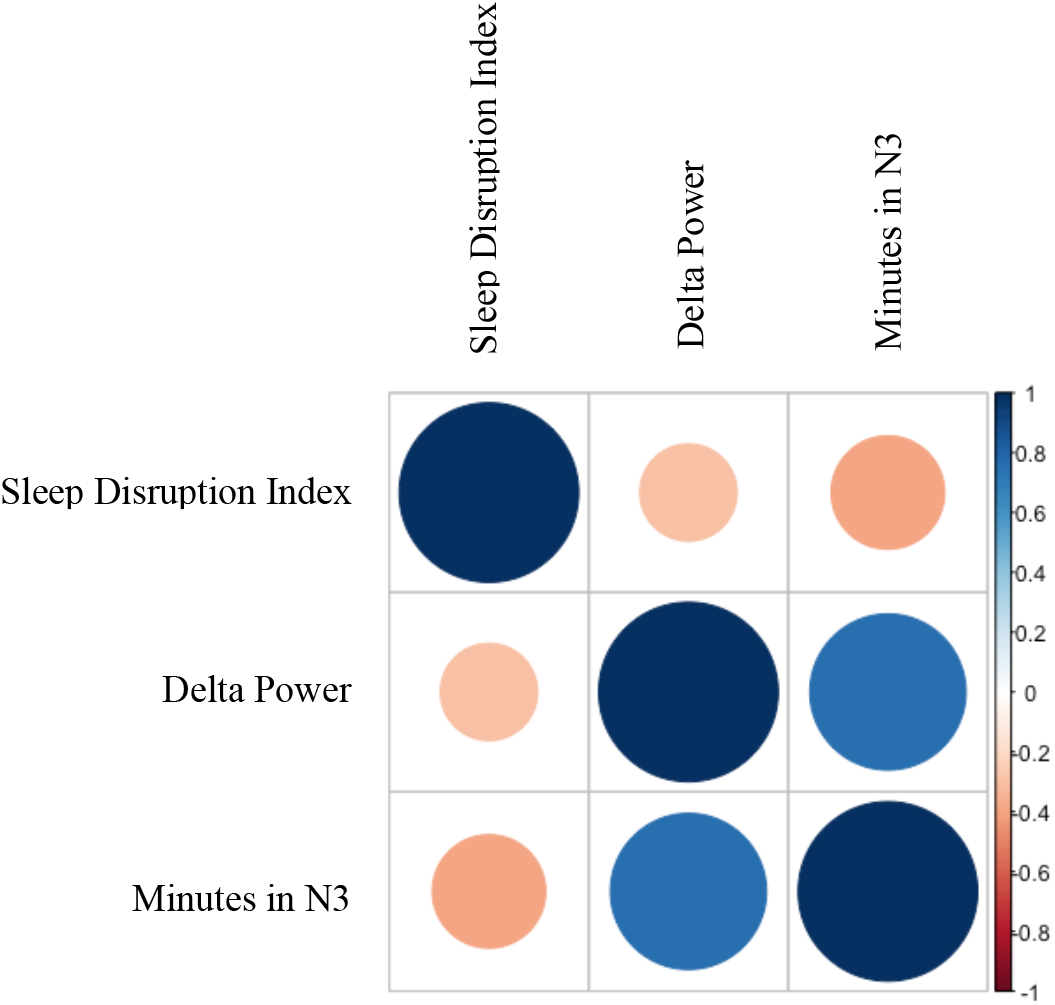
Correlation matrix of predictors (time spent in N3, delta power, and sleep disruption index) to be used in the multiple regression analysis predicting cueing benefit for response time and accuracy.

### Discussion

This experiment was designed to probe the effect of sleep reactivation on multiplication learning. Participants completed three blocks of six classes of multiplication problems with the groups coming from the six multiplicands 13, 14, 16, 17, 18, and 19. Each class was randomly paired with one of six sounds, before an afternoon nap with the final block serving as a Pre-test score. Increasing accuracy and decreasing response time over the three blocks indicate that participants did learn sufficiently. During the nap, three of the six previously learned sounds were presented in stage N2 and N3. After the nap, participants completed each of the 30 multiplication problems to serve as a Post-test score. Overall, participants in this group had a similar accuracy score before and after sleep while response time was slightly slower after the nap. However, there was no evidence that memory reactivation during a nap affected accuracy or response time as performance was similar for cued and uncued multiplication problems. Finally, after looking at performance by problem set for all 40 subjects, we were able to determine that based on accuracy and response time, problem sets 13, 14, and 16 seemed to be somewhat easier than problem sets 17, 18, and 19. This analysis can provide some input as to how easier and harder problems were differentially effected by TMR.

There are a few possible explanations for these findings. First, sounds may have been too soft to be operative during the nap. Our protocol to guard against this was to increase sound volume after 10 rounds of cueing, and we also decreased it if scalp EEG indicated signs of arousal. Because these precautions were taken to avoid playing sounds too softly throughout the nap, it is unlikely that this contributed greatly to the lack of cueing benefit found. We later explored the post-hoc hypothesis that the cues presented during sleep negatively influenced the possible TMR benefit through sleep disruption. It was determined there was no significant relationship between cueing benefit and sleep disruption index (Figure 6). Furthermore, the majority of participants had a low sleep disruption index score indicating that on average, cues during the nap were not disruptive.

Another possibility is that learning prior to sleep was inadequate, such that arbitrary sounds were not fully associated with classes of multiplication problems. To look into this possibility, we examined accuracy on tests of sound-problem associations. We found that participants averaged 79% correct and 19 participants achieved a perfect score on at least 1 of 5 sound tests administered. These findings argue against the idea that the associations were not well learned. It could be, however, that the same sound-problem practice may not have been maintained during sleep, which would have resulted in no benefit for cued items.

Next, it is possible that the paradigm may benefit from reactivation in a different sleep stage. In this study sounds were cued during stage N2 and N3, as deeper stages have been linked with greater memory performance and previous TMR studies have had success cueing in these stages. Though it is more difficult to target cueing to stage N1 because of the possibility of inducing an arousal, a follow-up study would be needed to assess this critique.

It is also a possibility that TMR benefits could have emerged after delayed testing. Because participants were only tested immediately after the nap, possible effects of delayed testing that would allow for more forgetting could have been missed. Such delayed TMR effects have been recently reported (Cairney, El Marj, & Staresina, 2018).

Another possibility is that multiplication learning reached a plateau such that memory reactivation could not move performance further. In other words, memory reactivation during sleep could be generally helpful for multiplication learning, but only when the learning begins to decline. With these learning and testing parameters, the skill knowledge gained may have stabilized. Perhaps at a further delay after learning, memory reactivation could act to offset a decline such that only those problems not reactivated would decline to a baseline level, presumably the level achieved when each problem was initially presented.

Finally, it may be that problem that different sets were not computed the same way. Fourteen participants reported solving multiplication problems in a strict, standard simple multiplication and addition fashion while others reported memorization, splitting the problems into smaller multiples, or a combination of the mentioned strategies. If participants used different strategies depending on the multiplier, TMR may not benefit those specifically cued during the nap but instead affect all problem sets. Likewise, TMR may have been absent if the cues promoted reactivation of multiplication learning generally, not in a way that favored the cued problems. To look into this final possibility, a follow-up wake study was conducted to allow us to assess the effect of sleep in this math learning paradigm.

## Experiment 3

### Methods

#### Participants

Nineteen undergraduate students (3 males, mean age = 21.00 ± 3.38 (SD)) consented to participate in this study and were paid for their time in the lab.

#### Procedure

Participants arrived at the lab between the hours of 11:00 am and 3:00 pm for Session 1. During this session, participants completed the same learning paradigm as those in Study 1. After learning, participants also completed a Pre-test. At this point, participants were allowed to leave the lab and returned two hours later for Session 2. Upon arrival, the Post-test was administered. Participants were instructed not to sleep during the break between Sessions 1 and 2 and were asked to describe what they did during this period.

## Results and Discussion

### Behavioral Analysis

Experiment 1 and 2 demonstrated that participants were able to learn over time with an average increase in accuracy and faster responses before sleep. Performance for learning before and after sleep is shown below in Figure 14.

**Figure 14.**
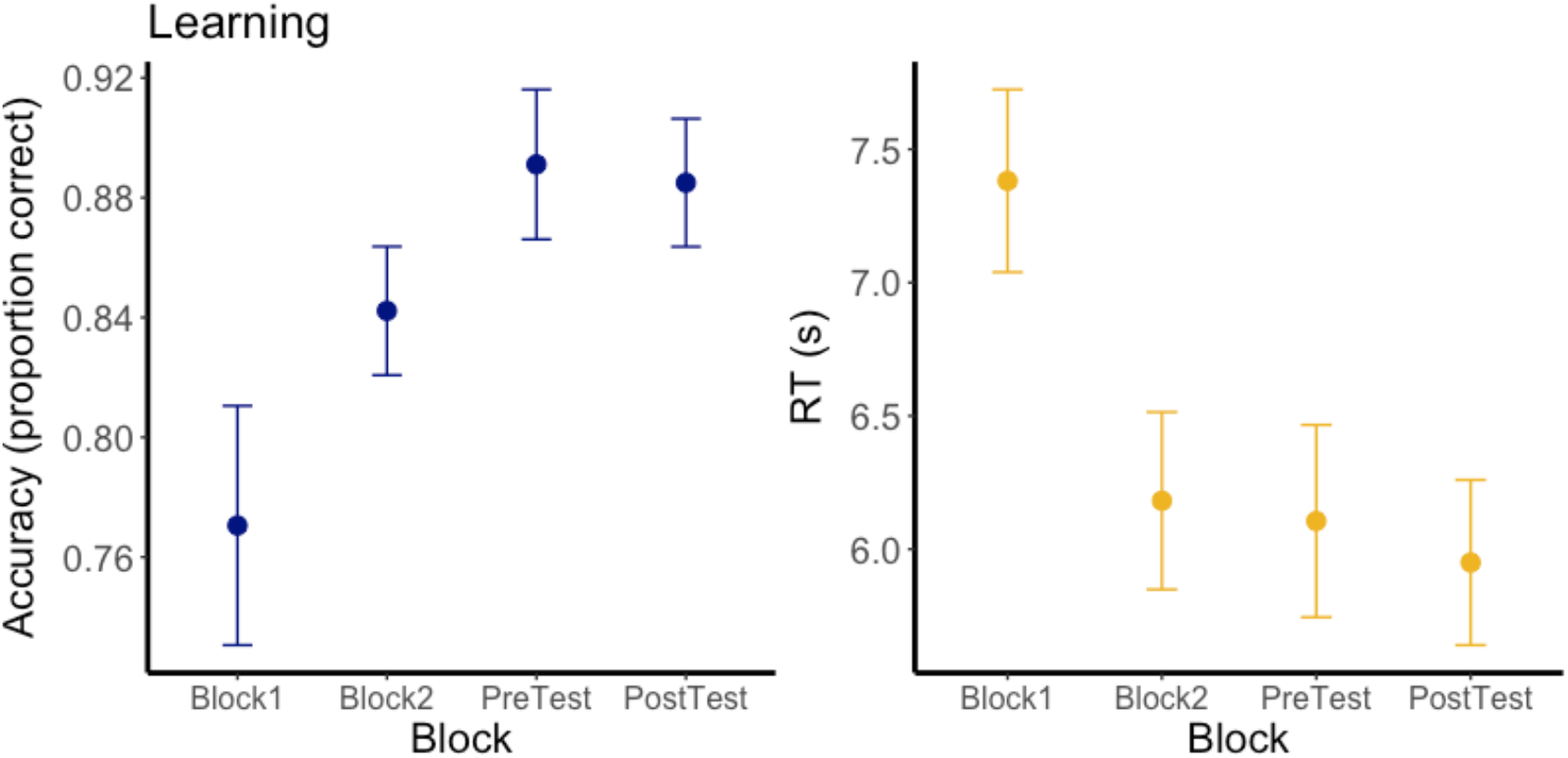
Performance for accuracy (left) and response time (right) throughout the experiment.

Behavior was compared for those who slept and those who did not between the Pre-test and Post-test. Participants in the nap group had an average accuracy of 0.88 ± 0.74 (SE) before sleep and 0.88 ± 0.66 (SE) after sleep. The participants who remained awake had an average accuracy of 0.89 ± 0.73 (SE) at the Pre-test and had a final accuracy of 0.89 ± 0.62 (SE). When looking at the response time, the nap group averaged 6.29 s ± 0.33 s (SE) for the Pre-test and 6.32 s ± 0.35s (SE) for the Post-test. The wake group had an average response time of 5.93 s ± 0.38 s (SE) and 5.79 s ± 0.33 s (SE) for the Pre-test and Post-test, respectively (Figure 15). A *t*-test revealed that performance for the two groups was matched for accuracy (*t* _(37.96)_ = 0.40, *p* = 0.69) and response time (*t* _(36.58)_ = 0.71, *p* = 0.48) at Pre-test.

**Figure 15.**
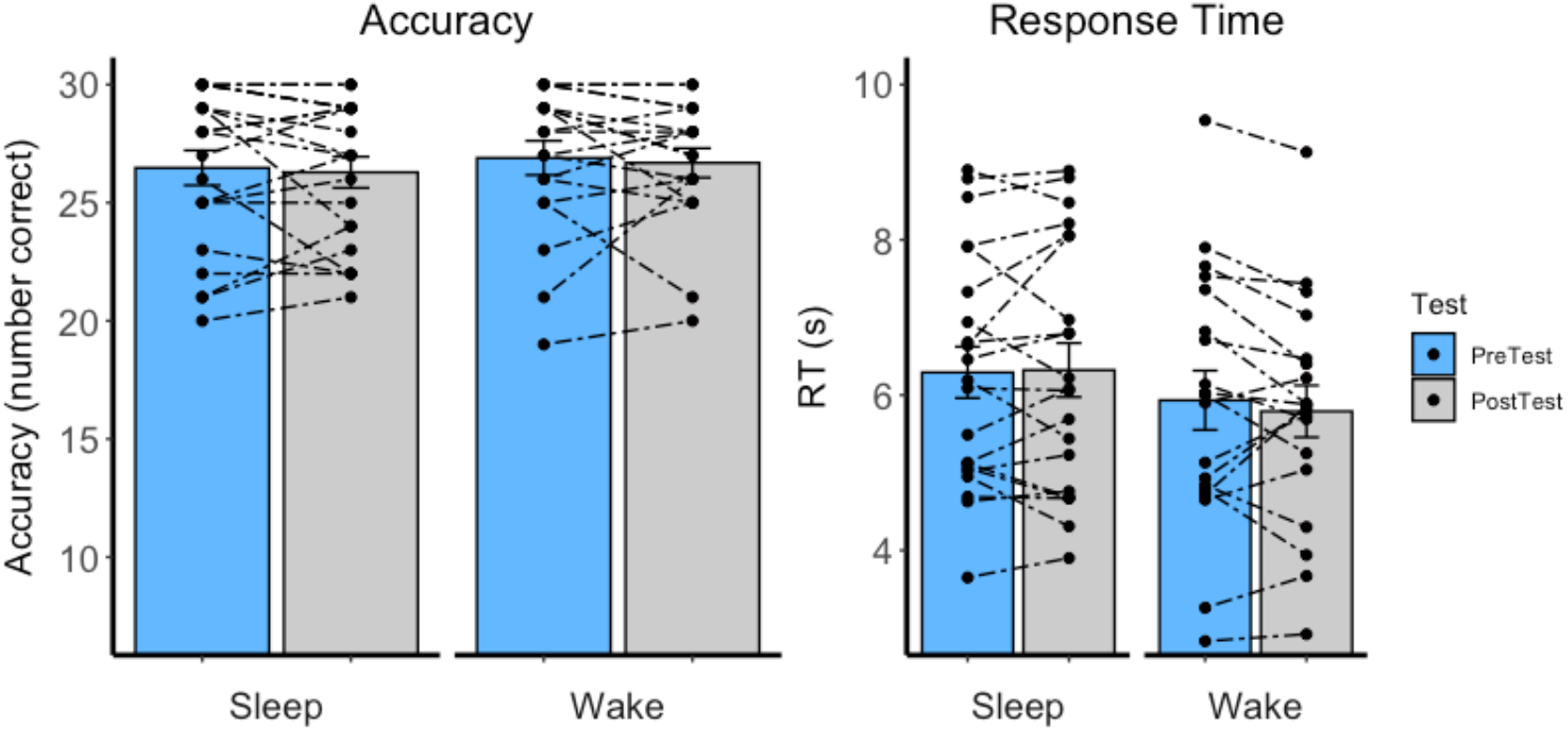
Pre-test and Post-test performance in accuracy (left) and response time (right) for Wake and Sleep subjects.

A 2 (Test: Pre-test/Post-test) x 2 (Group: Wake/Sleep) repeated measures ANOVA was conducted to analyze accuracy and response time. For accuracy, there was no main effect of Test (*F*_(1, 38)_ = 0.39, *p* = 0.54), Group (*F*_(1, 38)_ = 0.20, *p* = 0.66), or Test x Group interaction (*F*_(1, 38)_ = 0.001, *p* = 0.98). Similarly, for response time, there was no main effect of Test (*F*_(1, 38)_ = 0.31, *p* = 0.58), Group (*F*_(1, 38)_ = 0.85, *p* = 0.36), or Test x Group interaction (*F*_(1, 38)_ = 0.84, *p* = 0.37).

We next examined the performance on transfer problems completed on the Post-test. Participants in this group had an average accuracy of 0.89 ± 0.02 (SE) and average response time of 5.46s ± 0.38 (SE). Because there was no difference in either performance measure both types of problem (cued and uncued) in the nap group, results were collapsed to create a sleep group. To compare performance for the sleep and wake groups, we conducted a two-sided *student’s* two-sample t-test. This analysis revealed a marginally significant effect for accuracy (*t*(38) = 1.94, *p* = 0.06) with the wake group having a higher accuracy for transfer problems at the Post-test. For response time, this analysis revealed no significant difference between the sleep and wake groups (*t*(38) = 1.22, *p* = 0.23) (Figure 16).

**Figure 16.**
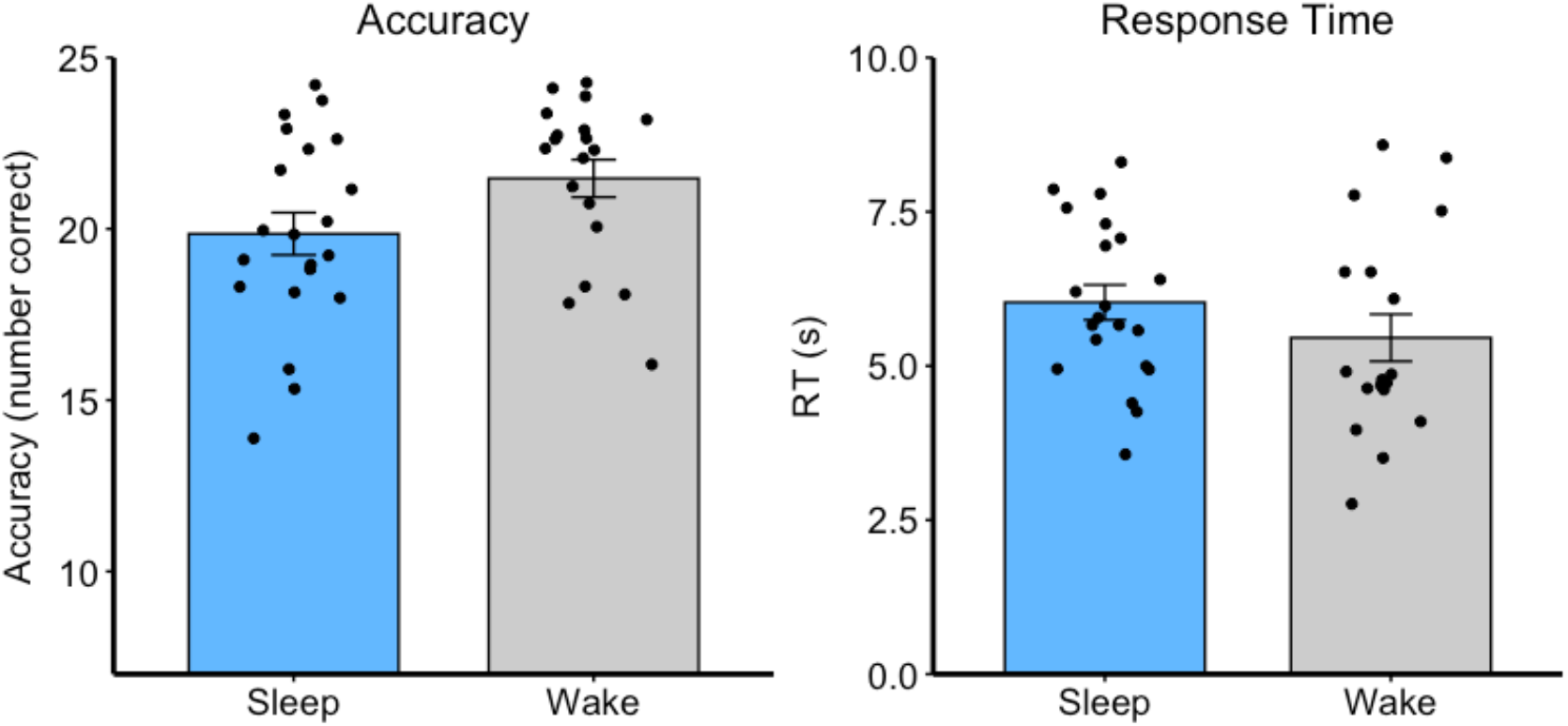
Performance for transfer problems completed at the Post-test. Accuracy (left) and response time (right) are depicted and separated by the Sleep and Wake groups.

### Discussion

We compared results from the participants who slept between the Pre-test and Post-test to allow us to understand the effect of sleep for mathematical problem solving. As performance for cued and uncued multiplication problems was similar in the Pre-test and Post-test for the sleep group, the results were combined for this analysis. Because a period of sleep has been found to benefit performance in a number of different learning paradigms, we expected that sleep would improve one’s ability to calculate problems efficiently and accurately. It was determined, however, that the wake and sleep groups performed similarly on the Pre-test and Post-test for both accuracy and response time.

There are a number of possible explanations for why sleep did not have an effect on performance for multiplication problems. First, it may be that a nap is not a sufficient period of rest. Although 90-minute naps have had success for other paradigms, it is possible that for multiplication learning, overnight sleep or multiple sessions of sleep might be necessary to produce measurable effects. It could also be that multiplication learning is not a sleep-dependent process. If this is true, it might be that periods of wake practice are sufficient to produce improvements in both accuracy and speed.

We also found that participants who remained awake during the period between the Pre-test and Post-test had a higher accuracy for the new, transfer problems introduced in the final test. While the conclusions that can be drawn from the transfer problems used are limited, it is possible that sleep and/or reactivation that occurred during sleep may have limited the ability to generalize previously learned information for new problems.

## General Discussion

These studies were designed in an attempt to implicate memory reactivation during sleep in multiplication learning. In Experiment 1, participants completed a number of multiplication problems to inform us on general learning and improvement over time on this task. In Experiment 2, participants were cued during slow-wave sleep with three of six sounds which were randomly paired with different multiplication classes. We found no evidence for an effect of TMR. Because it is possible that reactivation during the nap may have affected more problems than those specifically targeted, Experiment 3 was completed to further examine the broader role of sleep in this paradigm. In this experiment, subjects completed the same Pre-test and Post-test procedure but remained awake between the two tests. It was determined that sleep did not have a differential effect on later performance compared to an equal period of time without sleep.

For Experiment 2, it is possible that although the arbitrary sound-problem pairings were learned, they did not serve as meaningful cues during the nap. If this is true, the memory of performing multiplication problems may not have been maintained and reactivated. It might also be that while memory reactivation for the multiplication problems did not produce immediate effects, improvements in performance may have been detectable if a test had been given a day or more later. Finally, it could be that different strategies used by participants may have affected memory reactivation. Based on written report, we can conclude that participants used a number of different strategies throughout the experiment and it is possible that TMR may have been effective in impacting performance for specific strategies over others. In Experiment 3, it was determined that a period of sleep did not affect performance after being compared to a period of wake. It might be that a 90-minute nap is not sufficient to produce sleep effects in this paradigm. Furthermore, it might be that multiplication learning is not a sleep-dependent process and that improvements can be achieved during wake practice.

Future experiments could assess other facets of the protocol used in this experiment. Because participants seemed to reach a plateau of performance, it might be worthwhile to decrease the level of learning before sleep to see if effects are seen after TMR or a delay as participants are given more room for improvement. Future experiments could also assess the importance of sound cues in this paradigm. Perhaps the randomly associated sounds did not serve as meaningful cues during sleep, and if so, exploring whether spoken word cues of the problem sets produce different effects could be informative. Further experiments that explore the use of different cues, strategy, delayed testing, and longer or increased periods of sleep in a multiplication learning paradigm might provide insight about the process of mathematical problem-solving.

## Conclusion

Computing different mathematical problems is a skill that many students learn to do quickly and accurately over time. Whereas sleep is considered important for many aspects of academic success and has been found to play an important role in many learning paradigms, it has not been studied with respect to the role of multiplication learning specifically. This experiment was completed to better understand the role of sleep and memory reactivation in a multiplication learning task. As we did not find evidence that memory reactivation or sleep influences multiplication learning in Experiments 2 and 3, the question remains open as to the role of reactivation and sleep in this context. Because mathematical problem-solving is necessary in an academic setting, further experiments should be done to understand the role of sleep and wake practice for learning.

